# Phosphorylated CheV interacts with a subset of chemoreceptors

**DOI:** 10.1101/2025.02.06.636884

**Authors:** Miguel A. Matilla, Mario Cano-Muñoz, Elizabet Monteagudo-Cascales, Tino Krell

## Abstract

Chemotaxis pathways are among the most complex signaling systems in bacteria. A central feature of these pathways is the ternary complex formed by chemoreceptors, the autokinase CheA, and the coupling proteins CheW and CheV. Whereas CheW is present in all chemotaxis pathways, CheV is primarily found in bacteria that contain many chemoreceptors. CheV is a fusion of a CheW-like domain to a phosphorylatable receiver domain. The roles of CheV and its phosphorylation are currently uncertain. *Pseudomonas aeruginosa* contains many chemoreceptors for which the cognate signals have been identified. Quantitative capillary chemotaxis assays of a *cheV* mutant revealed that responses to certain chemoeffectors, such as nitrate and α-ketoglutarate, were drastically reduced, while responses to others, such as amino acids and inorganic phosphate, were comparable to the wild type, indicating that CheV selectively acts on specific chemoreceptors. To study the mechanism of CheV action, we conducted protein-protein interaction experiments using isothermal titration calorimetry. These studies showed that unphosphorylated CheV fails to bind to cytosolic fragments of the McpN and PctA chemoreceptors, which mediate responses to nitrate and amino acids, respectively. In contrast, the phosphorylation-mimic CheV D238E bound with very high affinity (*K*_D_ = 8 nM) to McpN but failed to interact with PctA. Thus, CheV in *P. aeruginosa* binds to some chemoreceptors but not to others in a phosphorylation-dependent manner. These results suggest that CheV is a regulatory protein that modulates signaling through specific chemoreceptors. CheV may thus facilitate the coordination of chemotaxis responses in complex, multi-chemoreceptor systems.

**Importance:** Of all chemosensory signaling proteins, CheV is perhaps the least understood. Our demonstration that CheV interacts only with certain chemoreceptors offers fundamental new insights. These findings, combined with the observation that CheV is present in bacteria with numerous chemoreceptors, suggest that CheV plays a role in coordinating chemotactic outputs in complex chemosensory systems. Understanding the mechanisms by which chemotactic responses are defined in bacteria with a high number of chemoreceptors is a major research priority in the field of chemotaxis. While previous studies, including this one, show that the ability to be phosphorylated is crucial for CheV function, the molecular consequences of CheV phosphorylation have remained unclear. Our discovery that phosphorylation is essential for CheV binding to certain chemoreceptors fills in this critical gap in understanding the molecular mechanism of CheV. This study is likely to inspire further research into CheV function in other bacteria using similar approaches.

## Introduction

Chemosensory pathways are among the most sophisticated signal transduction systems in bacteria (*1*, *2*). Genome analyses reveal that more than half of the sequenced bacterial genomes encode chemosensory signaling proteins (*3*). While most chemosensory pathways mediate chemotaxis, others carry out alternative cellular functions, like the control of second messenger levels, or are involved in twitching motility and mechanosensing (*3–5*).

The core of the chemosensory pathway is formed by a ternary complex comprised of chemoreceptors, the CheA autokinase, and a coupling protein. Signaling is typically initiated by ligand binding to the chemoreceptor ligand-binding domain (LBD), which in turn modulates the activity of CheA and the subsequent transfer of the phosphoryl group to the CheY response regulator. Other essential components of the pathway are the CheR methyltransferase and the CheB methylesterase, whose coordinate activities control the methylation state of the chemoreceptors (*1*, *2*).

Importantly, there are two coupling proteins in some bacteria, CheW and CheV (*6*, *7*). CheW consists of a single domain and is essential for the formation of hexagonally arranged chemosensory arrays (*8*). In contrast, CheV is a fusion of a CheW-like domain with a phosphorylatable receiver domain. All bacterial chemosensory pathways contain either CheW, CheV, or both (*7*, *9*). Several studies revealed a partial functional redundancy of CheW and CheV. Single deletions of the *cheV* or *cheW* genes in *Bacillus subtilis* (*10*) and *Campylobacter jejuni* (*11*) cause either no or only minor reductions in chemotaxis. However, the *cheV*/*cheW* double mutant of *B. subtilis* is completely defective for chemotaxis (*10*). Studies with other bacteria found a significant reduction, but not an absence of chemotaxis in *cheV* and *cheW* single mutants (*12–14*).

The phosphorylation of CheV is required for its activity (*15*). Experimental data and comparative genomic analyses support the idea that CheV acts as a phosphate sink that modulates the half-life of CheY-P (*16*, *17*). However, the functional consequences of CheV phosphorylation are unknown. Among all chemosensory signaling proteins, CheV is probably the least understood (*3*). However, interesting insight was provided by a bioinformatic analysis of *cheV* genes in enterobacteria (*16*). Bacteria lacking *cheV* tend to possess relatively few chemoreceptors (on average 5), whereas strains with *cheV* typically have a much higher number of chemoreceptors (on average 23). Furthermore, evolutionary analyses showed that CheV co-evolved with a specific family of chemoreceptors (*16*), suggesting that CheV may contribute differentially to chemotaxis mediated by different chemoreceptors. However, experimental data to support this hypothesis are lacking.

To address this question, we conducted quantitative chemotaxis experiments with *Pseudomonas aeruginosa* PAO1, a model strain in the study of chemotaxis (*18*). *P. aeruginosa* is among the most important human pathogens; it kills about half a million people annually (*19*, *20*). PAO1 has 5 gene clusters that encode chemosensory signaling proteins. These function in four distinct chemosensory pathways (Fig. 1) (*21*).

**Fig. 1.**
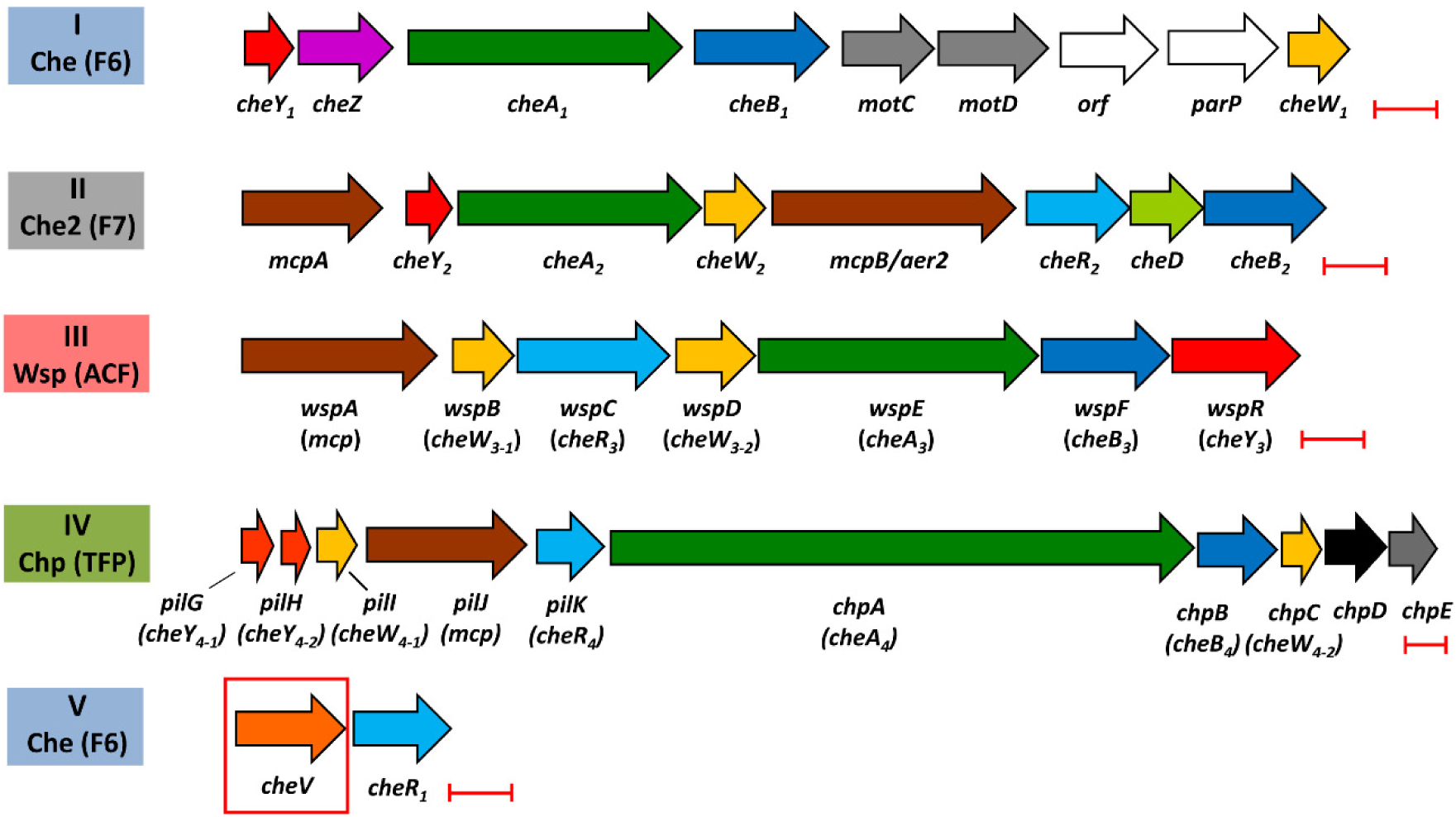
The five gene clusters that encode the signaling proteins of the four chemosensory pathways in *Pseudomonas aeruginosa* PAO1. Clusters I and V encode the proteins of the Che pathway (chemotaxis), cluster II encodes the Che2 pathway proteins (virulence by unknown mechanisms), cluster III corresponds to the Wsp pathway (c-di-GMP homeostasis), and cluster IV encodes the Chp pathway (twitching and mechanosensing). Pathway classification according to (*3*) is shown. Wsp: wrinkly spreader phenotype; Chp: chemosensory pili; F: flagellar motility; ACF: alternative cellular functions; TFP: type IV pili. Scale bars: 0.5 kbp.

The Che pathway mediates chemotaxis, the Che2 pathway plays a role in virulence by an unknown mechanism, the Wsp pathway regulates c-di-GMP levels, and the Chp system is involved in twitching and mechanosensing (*5*, *21–23*). Signaling through all four pathways is important for efficient virulence (*21*).

Strain PAO1 has 26 chemoreceptors. Bioinformatic and experimental data suggest that 23 receptors stimulate the Che pathway, whereas each of the remaining three chemoreceptors feeds into either the Che2, Wsp, or Chp pathway (*24*). PAO1 has a single CheV that is encoded in chemotaxis gene cluster V (Fig. 1). The signals recognized by about half of the PAO1 chemoreceptors have been identified (*25*). In most cases, mutants defective in these chemoreceptors showed either a complete loss or a large reduction in the chemotactic responses to their cognate ligands, indicating that the corresponding wild-type (wt) responses are primarily due to the action of a single chemoreceptor (see Table 1). This, and the wealth of information available on the function of its chemoreceptors makes PAO1 a valuable system well-suited to assess the contribution of CheV to the responses mediated by specific receptors.

**Table 1.** Summary of data available on chemoreceptors that mediate chemotactic responses to different chemoeffectors in *P. aeruginosa* PAO1. Responses of the wt strains and the *cheV* mutant are shown in Fig. 3. LBD families are defined according to Pfam (*66*) and, in the absence of annotation, by inspection of the Alphafold2 (*67*) model. Chemotactic responses that were significantly different in the *cheV* mutant compared to the wt strain are shown in boldface.

| Chemoeffector | Receptor | Receptor LBD family | Ref.<sup>a</sup> | Chemotaxis in *cheV* mutant relative to wt (%) |
| --- | --- | --- | --- | --- |
| **NaNO<sub>3</sub>** | **McpN** | **PilJ** | (28) | **5** |
| L-malate | CtpM | sCache_2 | (44, 68) | 92 |
| **Histamine** | **TlpQ** | **dCache<sup>b</sup>** | (46) | **51** |
| **α-ketoglutarate** | **McpK** | **HBM** | (42) | **23** |
| Inorganic phosphate | CtpL<br>CtpH | HBM<sup>b</sup><br>4HB<sup>b</sup> | (45, 69) | 39<br>94 |
| L-Val | PctA | dCache_1 | (29, 70, 71) | 83 |
| L-Gln | PctB | dCache_1 | (29, 70, 71) | 100 |
| γ-aminobutyrate | PctC | dCache_1 | (29, 71, 72) | 101 |
| **Acetylcholine** | **PctD** | **dCache<sup>b</sup>** | (43) | **45** |
| **Oxygen** | **Aer/Aer2** | **PAS** | (47) |  |

<sup>a</sup> References that document the functional relationship between chemoreceptor and chemoeffector.
<sup>b</sup> Inspection of an AlphaFold2 model.

We show that CheV participates in the signaling of only a subset of chemoreceptors, highlighting its role in regulating complex responses in systems with many chemoreceptors. We also demonstrate the functional relevance of CheV phosphorylation by showing that only a CheV phospho-mimic, but not unphosphorylated CheV, binds to chemoreceptors. Our study provides novel insight into the physiological function of one of the least understood chemosensory signaling proteins. It should provide an impetus to explore the role of CheV in other bacteria.

## Results

### CheW_1_ of *P. aeruginosa* is required for chemotaxis

*P. aeruginosa* PAO1 has 6 CheW homologs (Fig. 1) (*9*). Gene clusters III (Wsp pathway) and IV (Chp pathway) each harbor two *cheW* genes, whereas there is a single *cheW* gene in clusters I (Che pathway) and II (Che2 pathway). These CheW proteins share a modest sequence identity of 14 to 30 % (Fig. S1, Table S1). We assessed the role of 5 CheW homologs associated with the four chemosensory pathways: CheW_1_, CheW_2_, WspB, WspD, and ChpC, and conducted quantitative capillary chemotaxis assays with the wt strain and single mutants in each of the respective genes encoding CheW homologs to 1 mM L-cysteine, which triggers a strong chemoattractant response in strain PAO1. We observed a lack of chemoattraction in the *cheW*_1_ mutant (Fig. 2), which corresponds to the *cheW* homolog in the Che pathway (Fig. 1). The responses of the remaining 4 mutants were either similar or superior to those of the wt (Fig. 2).

**Fig. 2.**
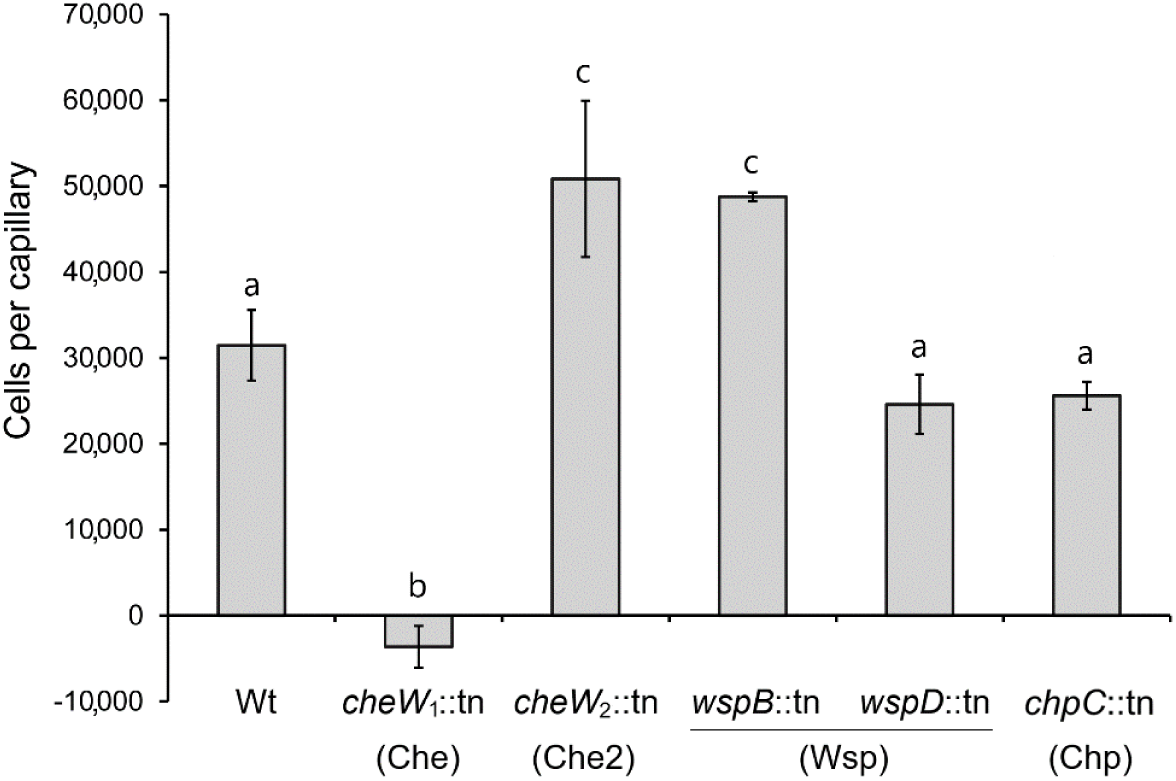
Specificity of interaction of CheW_1_ with the Che pathway. Quantitative capillary chemotaxis assays of *P. aeruginosa* PAO1 and five *cheW* mutants. The attractant was 1 mM L-cysteine. Data were corrected for the number of bacteria that swam into buffer-containing capillaries (14,400 ± 901 for wt; 13,800 ± 3,400 for *cheW*_1_::tn; 18,444 ± 4,986 for *cheW*_2_::tn; 17,333 ± 2,142 for *wspB*::tn; 16,222 ± 4,384 for *wspD*::tn; 13,200 ± 1,047 for *chpC*::tn). The corresponding chemosensory pathways are indicated in brackets. Bars with the same letter are not significantly different (P-value < 0.05; by Student’s t-test).

These findings indicate a specific interaction of the CheW encoded in the Che gene cluster with the other proteins of the Che pathway, supporting previous findings that the *P. aeruginosa* chemosensory proteins assemble into insulated pathways (*23*, *26*).

### CheV is required for chemotaxis mediated by a subset of *P. aeruginosa* chemoreceptors

*P. aeruginosa* has one CheV protein, which is encoded in the chemosensory gene cluster V together with the CheR_1_ methyltransferase (Fig. 1) (9, 21, 24). To assess the contribution of CheV to responses mediated by different chemoreceptors, we selected chemoeffectors for which the corresponding chemoreceptor has been identified (Table 1). In most of the cases, single chemoreceptor mutants are unable to mediate chemotaxis to their cognate ligands (Table 1). This permits us to attribute a specific response to individual chemoreceptors.

Quantitative capillary chemotaxis assays to four chemoeffectors recognized by McpN, McpK, TlpQ and PctD, respectively, and aerotaxis responses were significantly reduced in the *cheV* mutant as compared to the wt strain (Fig. 3 A, B).

**Fig. 3.**
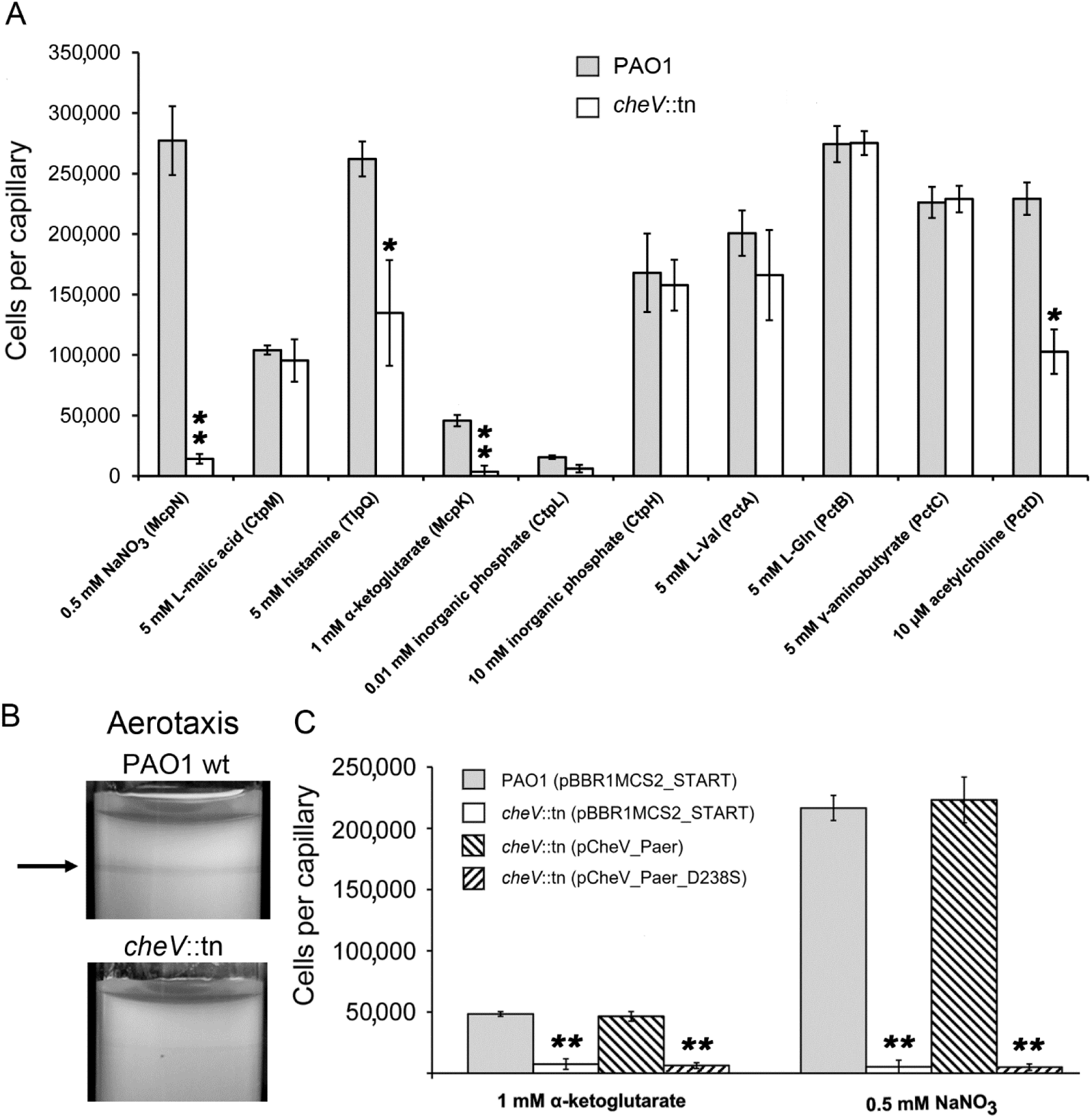
Tactic responses of *P. aeruginosa* and its *cheV* mutant towards different signals. **A)** Quantitative capillary chemotaxis assays to the indicated compounds. The corresponding chemoreceptors are shown in brackets. Data were corrected for the number of bacteria that swam into buffer-containing capillaries (14,130 ± 2,232 for wt; 11,366 ± 2,221 for *cheV*::tn). Information on the chemoreceptor-chemoeffector interaction is provided in Table 1. **B)** Aerotaxis tube assays. The arrow points to the band characteristic for an aerotaxis response. **C)** Quantitative capillary chemotaxis assays to α-ketoglutarate and nitrate of *P. aeruginosa* strains harboring different pBBR-based plasmids. Data were corrected for the number of bacteria that swam into buffer-containing capillaries (8,725 ± 3,344 for PAO1 (pBBR1MCS2_START); 6,208 ± 2,734 for *cheV*::tn (pBBR1MCS2_START); 9,883 ± 3,514 for *cheV*::tn (pCheV_Paer); 8,291 ± 2,227 for *cheV*::tn (pCheV_Paer_D238S). pBBR1MCS2_START: empty plasmid; pCheV_Paer: pBBR1MCS2_START plasmid encoding wt CheV; pCheV_Paer_D238S: pBBR1MCS2_START plasmid encoding the CheV D238S variant, in which the phosphoryl-accepting Asp residue has been replaced by Ser. Student’s t-test: *P-value <0.05, ** P-value<0.01.

Chemotaxis to the remaining 6 chemoeffectors tested was comparable to that of the wt strain (Fig. 3A). Responses to nitrate and α-ketoglutarate were down by ∼95 and ∼92 %, respectively, and chemotaxis to histamine and acetylcholine were reduced by about half in the *cheV* mutant (Fig. 3A). Aerotaxis assays of the wt strain showed the characteristic band for aerotaxis close to the air interface. This band was not obvious with the *cheV* mutant (Fig. 3B), indicating that CheV is important for aerotaxis. The chemoreceptors whose responses were diminished in the *cheV* mutant belong to different families containing PilJ, dCache, HBM or PAS-type LBDs (Table 1). Furthermore, the aerotaxis receptor Aer contains a cytosolic LBD, whereas the LBDs of the remaining chemoreceptors are located in the periplasm. The signaling domains of all receptors belong to the 40 H (heptad repeat) family (*9*, *27*).

When *cheV* was expressed *in trans* in the *cheV*-deficient strain, chemotaxis to α-ketoglutarate and nitrate in the *cheV* mutant was fully restored (Fig. 3C). However, no restoration of chemotaxis was observed when the plasmid encoded CheV in which the phosphoryl-group-accepting aspartate (D238) was replaced with serine (Fig. 3C). Thus, phosphorylation was essential for CheV activity. This finding is consistent with previous studies on *B. subtilis* CheV (*15*).

### CheV and CheW_1_ are both required for chemotaxis to nitrate and α-ketoglutarate

As shown above, inactivation of *cheV* had the strongest effect on chemotaxis to nitrate and α-ketoglutarate (Fig. 3A). To assess the role of CheW_1_ in these responses, we conducted chemotaxis assays with the *cheW*_1_ mutant. Like the *cheV* mutant, the *cheW*_1_ mutant failed to respond to nitrate and α-ketoglutarate (Fig. 4). This result highlights the essential roles of CheW_1_ and CheV in mediating these responses and indicates that CheW and CheV are not functionally redundant, as has been suggested (*10*, *11*).

**Fig. 4.**
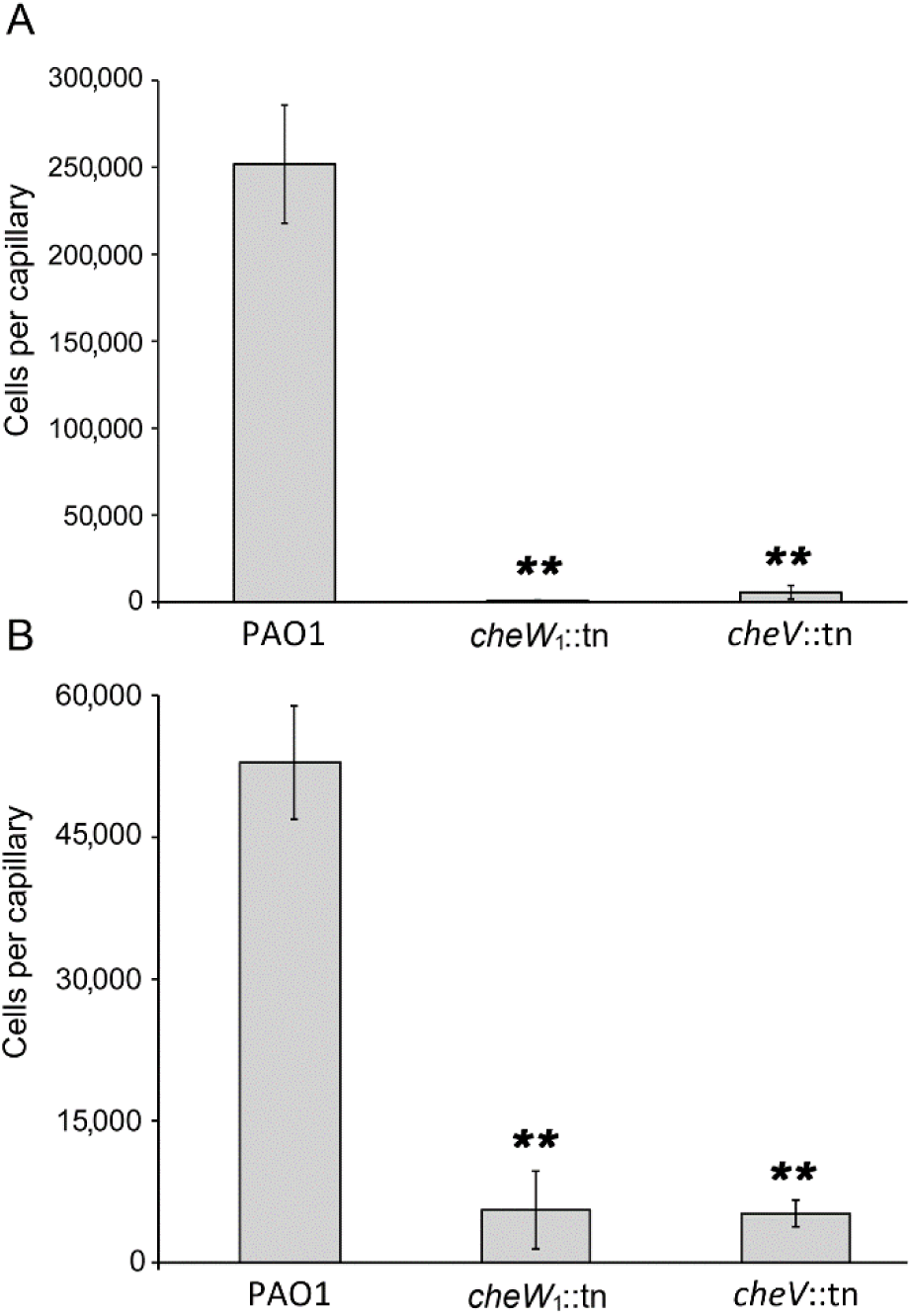
Both CheW_1_ and CheV are required for chemotactic responses of *P. aeruginosa* PAO1 to nitrate and α-ketoglutarate. Quantitative capillary chemotaxis assays of the wt and mutants in *cheW*_1_ or *cheV* to 0.5 mM NaNO_3_ (**A**) or 1 mM α-ketoglutarate (**B**). Data were corrected for the number of bacteria that swam into buffer-containing capillaries (20,227 ± 2,314 for PAO1 wt; 9,911 ± 3,694 for *cheW*_1_::tn; 13,050 ± 4,340 for *cheV*::tn). Student’s t-test: ** P-value<0.01.

### A phosphorylation mimic of CheV binds to a cytosolic fragment of McpN but not to a cytosolic fragment of PctA

The *P. aeruginosa cheV* mutant fails to respond to nitrate, whereas responses to different amino acids were comparable to the wt strain (Fig. 3A). Nitrate chemotaxis is mediated by the McpN chemoreceptor (*28*), whereas amino acid chemotaxis is mediated by the three receptors PctA, PctB and PctC (*29*). The latter three receptors are paralogous, and their cytosolic fragments share 93 % sequence identity (*30*). To test whether CheV interacts with McpN but not with PctA, we generated pET-based expression vectors containing the *cheV* gene and the DNA sequences encoding the cytosolic fragments of McpN (McpN_CF) and PctA (PctA_CF). The corresponding proteins were overexpressed in *E. coli* and purified. We then conducted isothermal titration calorimetry (ITC) experiments to study protein-protein interactions.

The control titration of buffer with CheV resulted in small and uniform peaks, representing dilution heats (Fig. 5A).

**Fig. 5.**
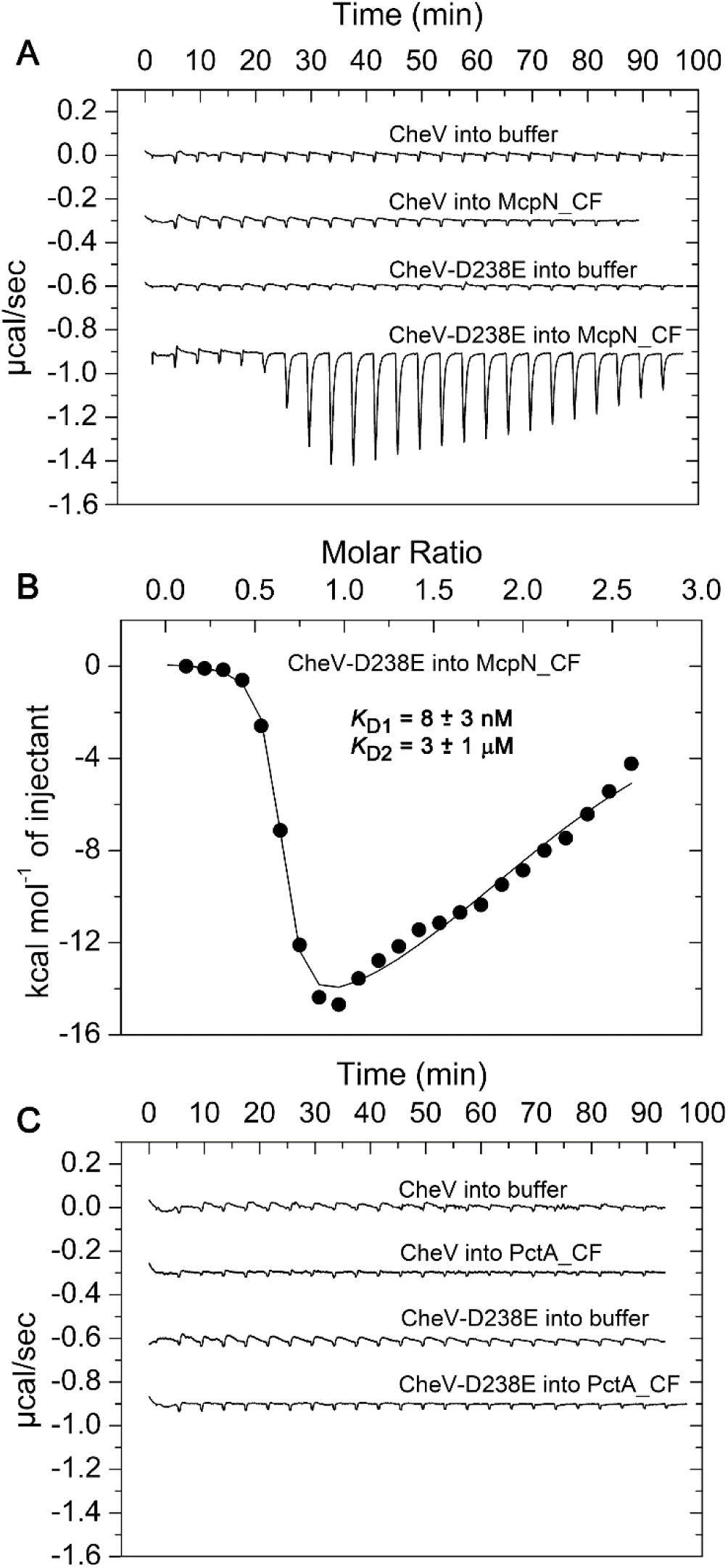
Isothermal titration calorimetry study of the binding of CheV and the phosphorylation mimic CheV D238E to the cytosolic fragments of the McpN and PctA chemoreceptors. Titration of either 10 µM McpN_CF (**A**) or PctA_CF (**C**) with 12.8 µl aliquots of 113 µM CheV or CheV-D238E. Shown are also the control titration of buffer with CheV and CheV-D238E. **B**) Concentration-normalized and dilution-heat-corrected titration data for the binding of CheV-D238E to McpN_CF. Data were fitted with the "Two Binding Sites" model in the MicroCal version of Origin software for ITC.

A similar curve was obtained when McpN_CF was titrated with unphosphorylated CheV. Since the phospho-Asp bond is labile, no stably phosphorylated receiver domains can be generated for experimentation (*31*). However, numerous studies show that the replacement of the phospho-accepting aspartate with glutamate mimics protein phosphorylation (*32–34*). Therefore, we generated the CheV D238E mutant for use in ITC assays. Titration of McpN_CF with CheV D238E resulted in a response in which two binding events could be distinguished: an initial high-affinity endothermic event with a *K*_D_ of 8 ± 3 nM followed by an exothermic event of lower affinity (*K*_D_= 3 ± 1 µM). An n-value of close to 0.5 was obtained for the first event indicative for the binding of a CheV D238E monomer to the McpN_CF dimer. No reliable information on the stoichiometry of interaction can be obtained from hyperbolic traces such as the second exothermic event. We hypothesize that this second event represents binding of another CheV to the opposing face of the McpN_CF dimer (Fig. S2). As shown in Fig. 5C, neither CheV nor CheV D238E bound to PctA_CF. The differential interaction of CheV D238E thus agrees with the failure of the *cheV* mutant to respond to nitrate while retaining responses mediated by PctA, PctB, and PctC (Fig. 3A). Our data indicate that phosphorylation of CheV is required to interact with chemoreceptors and confirm the functional relevance of CheV phosphorylation.

## Discussion

The hexagonal chemosensory array is formed by chemoreceptors, the CheA autokinase, and at least one coupling protein. CheW is a coupling protein in all species, but many species also have CheV. The composition of these arrays is highly variable among bacteria (*35*) and may include additional proteins, as in *Vibrio cholera* (*36*, *37*). The coupling proteins are essential for the activity of chemosensory pathways. However, the importance of CheV is poorly understood, despite its being present in ∼60% of prokaryotes with chemotaxis systems (*16*). Also, the functional consequences of CheV phosphorylation have been unclear. However, it is noteworthy that CheV is typically present in bacteria that have many different chemoreceptors (*16*).

Previous studies have shown that: 1) CheV increases the kinase activity of CheA (*38*); 2) CheV is phosphorylated by CheA (*15*, *17*); 3) CheW and CheV integrate similarly into the chemoreceptor baseplate, indicating that both play a role in establishing the array structure (*35*, *39*); 4) the interaction between CheV and chemoreceptors occurs through the CheW-like domain of CheV (*11*); and 5) CheV has co-evolved with certain chemoreceptors (*16*). We show here that, in *Pseudomonas aeruginosa*: 1) CheV participates in signaling through the McpN and McpK chemoreceptors (Fig. 3A); 2) both CheV and CheW are essential for signaling through these two chemoreceptors (Fig. 4); 3) the non-phosphorylatable D238S variant of CheV is inactive (Fig. 3C); 4) unphosphorylated CheV fails to bind to the cytosolic fragments of chemoreceptors (Fig. 5); and 5) the phosphorylation mimic D238E variant of CheV binds with high affinity to the cytosolic fragment of McpN, but not to the cytosolic fragment of PctA (Fig. 5).

These results suggest that CheV is a regulatory protein that modulates signaling through specific chemoreceptors (Fig. 6).

**Fig. 6.**
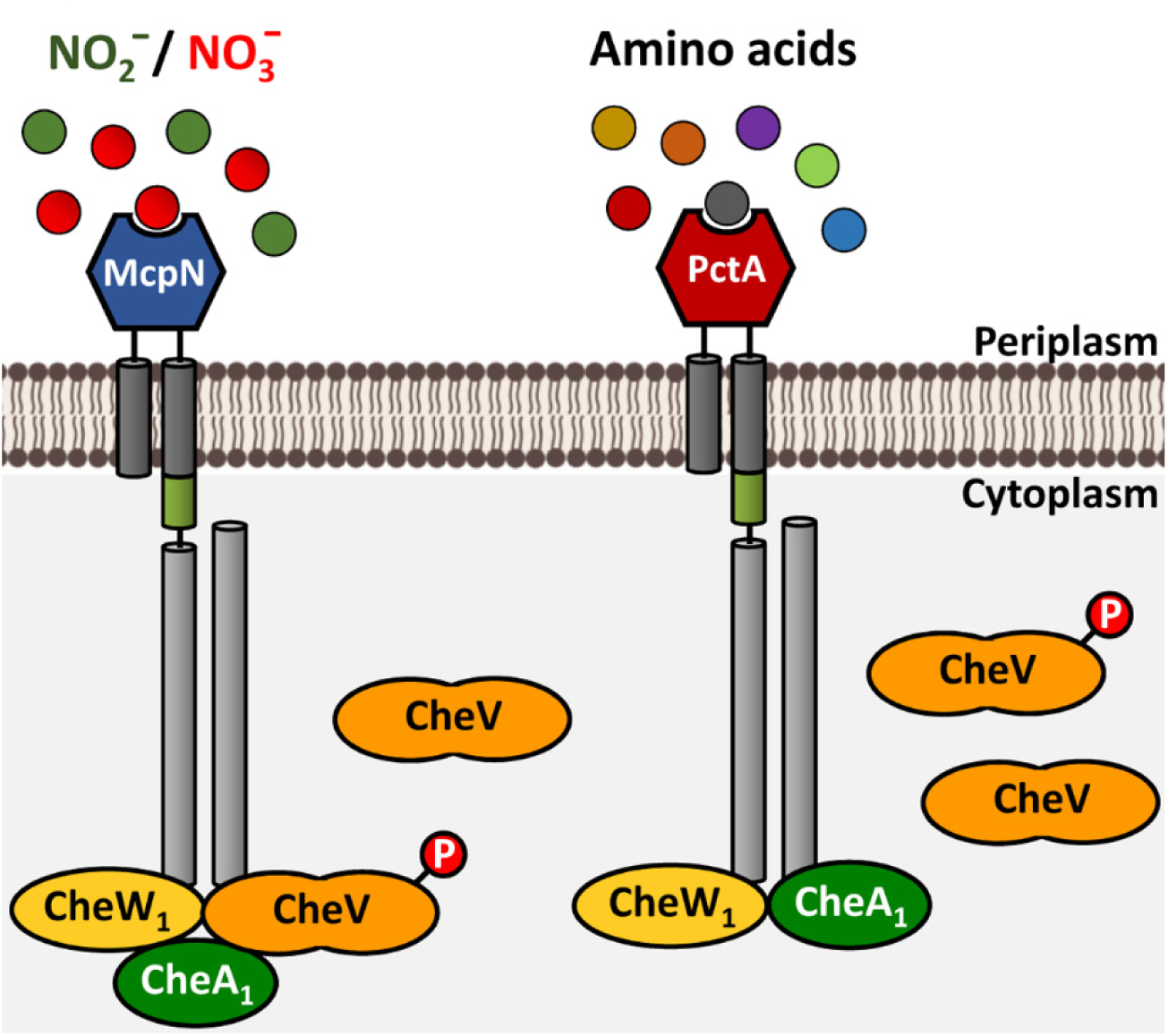
Schematic illustration of the differential roles of CheV in mediating *Pseudomonas aeruginosa* chemotaxis. The chemoreceptor McpN (CheV sensitive) mediates responses to nitrate and nitrite, whereas PctA (CheV insensitive) is a chemoreceptor for a broad-range of amino acids. Only the phosphorylated form of CheV binds to the cytosolic domain of McpN.

Bacteria encoding CheV proteins have, on average, about five times more chemoreceptors than those without CheVs (*16*). Thus, CheV may facilitate the coordination of chemotaxis responses in complex, multi-chemoreceptor systems.

Our previous proteomics studies have reported the levels of chemosensory proteins in *P. aeruginosa* PAO1 grown under three different conditions (*40*, *41*). The intensity-based absolute quantification (iBAQ) values of CheV and the 6 versions of CheW are provided in Table S2. Several conclusions can be drawn from these data. 1) CheW_1_ and CheV are by far the most abundant coupling proteins, 2) There are only moderate differences in CheW_1_ and CheV levels during growth under different conditions. 3) Under all three growth conditions there was a 2 to 4-fold excess of CheW_1_ over CheV, which is consistent with the notion that CheV acts only on a subset of chemoreceptors.

Our data indicate that phosphorylation of CheV is required for interaction with McpN and, potentially, for the formation of the signaling complex. We hypothesize that the phosphorylation state of CheV may determine the magnitude of signaling through McpN and, by inference, McpK. In the absence of signal, CheA is in the “kinase on state,” and signal binding to the chemoreceptors decreases CheA kinase activity and thereby the concentration of CheY-P. Phosphorylated CheV would further decrease CheA activity to promote chemotaxis. This mechanism is consistent with the proposal that CheV may act as a phosphate sink (*16*).

We show that the inactivation of *cheV* significantly decreased chemotaxis to nitrate, histamine, α-ketoglutarate, acetylcholine, and oxygen, whereas responses to L-malic acid, inorganic phosphate, L-Val, L-Gln, and γ-aminobutyrate were unaffected (Fig. 3A). The responses to nitrate, α-ketoglutarate, and acetylcholine are each mediated by a single chemoreceptor: McpN (*28*), McpK (*42*) and PctD (*43*), respectively. Receptors that were insensitive to the presence of CheV were the malate receptor CtpM (*44*), the phosphate specific CtpL and CtpH receptors (*45*), and the three paralogous amino acid receptors PctA, PctB and PctC (*29*). Histamine chemotaxis was decreased but not eliminated in the absence of CheV. TlpQ is the primary receptor for histamine, but two other chemoreceptors also mediate histamine chemotaxis (*46*). If one or two of these receptors do not require CheV, this partial inhibition is explained. Similarly, there are two oxygen chemoreceptors (*47*).

What features distinguish thus CheV-dependent from CheV-independent receptors? There is no obvious correlation between a sequence clustering analysis of the cytosolic PAO1 chemoreceptor fragments and receptor dependence on CheV (Fig. S3). A number of studies have defined the CheW-binding site on chemoreceptors (*48–52*). Although the binding site for CheV has not been rigorously determined, some data suggest that it binds to the same site as CheW (*7*, *16*, *53*). A sequence alignment of McpN_CF (CheV-sensitive) with PctA_CF (CheV-insensitive) shows that the amino acids assigned to the CheW-binding site are conserved between both proteins (Fig. S4). Because CheV co-evolved with a particular chemoreceptor family, the CheW-binding site and phospho-CheV binding site are unlikely be identical (*16*). The critical differences may lie in regions located rather far from the known CheW-binding site.

A major gap in our knowledge about chemosensory pathways is in understanding to what degree, in bacteria with multiple chemosensory systems, these pathways cross-talk with each other and other signaling networks. Whereas some reports show that these pathways are insulated (*23*, *54*, *55*), other studies show that they communicate with other systems (*56–58*). The question of insularity is related to the degree of specificity with which the different signaling proteins assemble to pathways. The *cheW*_1_ mutant is largely impaired in chemotaxis (Fig. 2), indicating that the remaining CheW homologs are unable to participate in this pathway. The interaction of CheW homologs only with their corresponding pathways is consistent with previous findings showing specific signaling protein interaction (*23*, *26*). Our demonstration that a single CheW is needed for chemotaxis in *P. aeruginosa* is similar to the situation in *Vibrio cholerae* and *Rhodobacter sphaeroides.* They possess three or two chemotaxis signaling pathways, respectively (*9*), but use only a single CheW for chemotaxis (*59*, *60*). However, the spirochaete *Borrelia burgdorferi* requires two of its three CheW proteins for chemotaxis (*61*).

CheV was first discovered in *Bacillus subtilis*. Initial studies suggested redundancy between CheV and CheW, as mutants deleted for either gene maintained chemotaxis, whereas the double mutant was non-chemotactic (*10*). A similar redundancy has been observed in *Campylobacter jejuni* (*11*). In contrast, our results show that CheV and CheW_1_ are non-redundant; mutants in either the *cheV* or *cheW*_1_ gene are defective in nitrate and α-ketoglutarate chemotaxis (Fig. 4). Our findings are reminiscent of those made with *Helicobacter pylori*, in which absence of *cheW* completely abolishes chemotaxis and the absence *cheV_1_* severely compromises chemotaxis (*38*, *53*, *62*). Our work provides the insight that CheV can interact with a subset of receptors to provide a mechanism for maintaining segregated control of their chemosensory signaling.

## Materials and methods

### Strains, plasmids, oligonucleotides and culture conditions

Bacteria and plasmids used in this study are described in Table S3, whereas oligonucleotides are listed in Table S4. *P. aeruginosa* strains were grown routinely at 30 °C and 37 °C, respectively, in lysogeny broth (LB) or M9 minimal medium supplemented with 6 mg/l Fe-citrate, trace elements (*63*) and 15 mM glucose as carbon source. *E. coli* strains were grown at 37 °C in LB. *E. coli* DH5α was used as a host for gene cloning. When appropriate, antibiotics were used at the following final concentrations (in μg/ml): ampicillin, 100; kanamycin, 50; streptomycin, 50 (*E. coli*) and 100 (*P. aeruginosa* strains); gentamicin, 10 (*E. coli* strains) and 50 (*P. aeruginosa* strains); tetracycline, 60; rifampin, 10; chloramphenicol, 25. Sucrose was added to a final concentration of 10 % (w/v) when required to select derivatives that had undergone a second crossover event during marker-exchange mutagenesis.

### In vitro nucleic acid techniques

Total DNA extraction was performed using the Wizard^®^ genomic DNA purification kit (Promega). Plasmid DNA was isolated using the NZY-Miniprep kit (NZY-Tech). For DNA digestion, alkaline phosphatase and ligation reactions, manufacturers’ instructions were followed (New England Biolabs and Roche). Competent cells were prepared using calcium chloride (*64*). Transformations and electroporations were performed following standard protocols (*64*). DNA fragments were recovered from agarose gels using the Qiagen gel extraction kit. PCRs were purified using the Qiagen PCR Clean-up kit. Phusion high-fidelity DNA polymerase (Thermo Fisher Scientific) was used in the amplification of PCR fragments for cloning. Sequences of these PCR fragments were verified by DNA sequencing.

### Construction of plasmids

For the construction of plasmids for protein overexpression and gene complementation, DNA sequences were amplified using primers described in Table S4 and cloned into pET28b(+) or pBBR-based plasmids, respectively. To generate pCheV_Paer_D238S and pET28b-CheV-D238E, the phosphorylatable aspartate (D238) of PA3349 (CheV) was replaced by serine or glutamic acid by overlapping PCR. Complementation plasmids were transformed into *P. aeruginosa* strains by electroporation.

### Chemotaxis assays

Overnight cultures in M9 minimal medium supplemented with 6 mg/l Fe-citrate, trace elements (*63*) and 15 mM glucose were used to inoculate fresh medium to an OD_660_ of 0.05. Cells were cultured at 37 °C (*P. aeruginosa*) to an OD_660_ of 0.4-0.5. Subsequently, cells were washed twice by centrifugation (1,667 x *g* for 5 min at room temperature) and resuspension in chemotaxis buffer (50 mM KH_2_PO_4_/K_2_HPO_4_, 20 mM EDTA, 0.05% [v/v] glycerol, pH 7.0). Cells were then resuspended in the same buffer at an OD_660_ of 0.1 and 230 µl aliquots of the resulting cell suspension were placed into the wells of 96-well microtiter plates. Then, 1-µl capillaries (Microcaps, Drummond Scientific) were heat-sealed at one end and filled with buffer (control) or chemoeffector solution prepared in chemotaxis buffer. The capillaries were rinsed with sterile water and immersed into the bacterial suspensions at their open ends. After 30 min, capillaries were removed from the wells, rinsed with sterile water, and emptied into 1 ml of chemotaxis buffer. Serial dilutions were plated onto M9 minimal medium plates supplemented with 15 mM glucose and incubated at 30 °C prior to colony counting. Data were corrected with the number of cells that swam into buffer containing capillaries. Data are the means and standard deviations of at least three biological replicates conducted in triplicate.

### Aerotaxis assays

These assays were carried out using the tube test method as described previously (*65*) with some modifications. Briefly, *P. aeruginosa* strains were grown in MS medium (30 mM Na_2_HPO_4,_ 20 mM KH_2_PO_4,_ 25 mM NH_4_NO_3,_ 1 mM MgSO_4_) supplemented with 6 mg/l Fe-citrate, trace elements (*63*) and 15 mM glucose. Fresh medium was inoculated to an OD_660_ of 0.05. At an OD_660_ of 0.4, cells were washed twice with chemotaxis buffer (50 mM KH_2_PO_4_/K_2_HPO_4_, 20 mM EDTA, 0.05% [v/v] glycerol, pH 7.0) and concentrated to an OD_660_ of 0.5 in the same buffer. Subsequently, 1.5 ml of cell suspensions were mixed with the same volume of 0.5 % (w/v) bacto-agar (Difco) prepared in chemotaxis buffer. This mixture was poured into sterile glass tubes and pictures were taken after 1 h incubation at 30 °C. Aerotactic responses are visualized by the formation of a clear band below the agarose/air interface.

### Protein overexpression and purification

*E. coli* BL21(DE3) harboring plasmids pET28b-CheV, pET28b-CheV-D238E, pET28b_McpN_CF and pET28b_PctA_CF were grown in 2-l Erlenmeyer flasks containing 500 ml LB medium supplemented with kanamycin. Cultures were grown under continuous stirring (200 rpm) at 30 °C. At an OD_660_ of 0.5-0.6, protein expression was induced by the addition of 0.1 mM isopropyl-β-D-thiogalactopyranoside. Growth was continued at 18 °C overnight and cells were harvested by centrifugation at 10,000 x *g* for 20 min at 4 °C. Proteins were purified by metal affinity chromatography. Cell pellets from a 1 l culture were resuspended in 40 ml of buffer A (30 mM Tris/HCL, 300 mM NaCl, 5 % (v/v) glycerol, 10 mM imidazole, 0.1 mM EDTA, 5 mM 2-mercaptoethanol, pH 8.5) containing 1 mM PMSF protease inhibitor (Thermo Fisher Scientific) and Benzonase (Merck). Cells were then broken by French press treatment at a gauge pressure of 1000 lb/in^2^. After centrifugation at 20,000 x *g* for 1 h, the supernatants were loaded onto a 5-mL HisTrap column (Amersham Bioscience) equilibrated in buffer A. After a wash with 40 ml of buffer A containing 40 mM imidazole, proteins were eluted by a linear gradient of 40 to 500 mM imidazole in buffer A.

### Isothermal titration calorimetry (ITC)

Titrations were performed on a VP microcalorimeter (MicroCal, Northampton, MA, USA) at 25 °C. Freshly purified proteins were dialyzed into 3 mM Tris, 3 mM PIPES, 3 mM MES, 5 mM 2-mercaptoethanol, pH 8.0. Ten µM solutions of either McpN_CF or PctA_CF were placed into the calorimeter sample cell and titrated at 240-second intervals with 12.8 µl aliquots of 113 µM CheV or CheV D238E. To correct for reactant dilution, the average enthalpy of the final peaks observed after saturation was subtracted from the titration data. The data were normalized to the ligand concentrations and the first data point removed. Data were analyzed using the "Two Binding Sites" model in the MicroCal version of Origin software for ITC.

## Acknowledgements

We are indebted to Michael Manson for editing the manuscript and providing constructive scientific comments. We thank Raquel Vázquez Santiago for technical support.

## Author contributions

MAM, MCM, and EMC conducted research and analysed data, TK and MAM conceived study and designed experiments, TK wrote initial draft of the manuscript, all authors edited the manuscript. All authors have seen an approve the final version of the manuscript.

## Data and materials availability

The datasets used and/or analyzed during the current study are available from the corresponding author on reasonable request.

## Supplementary Information

### Supplementary Figures

**Fig. S1.**
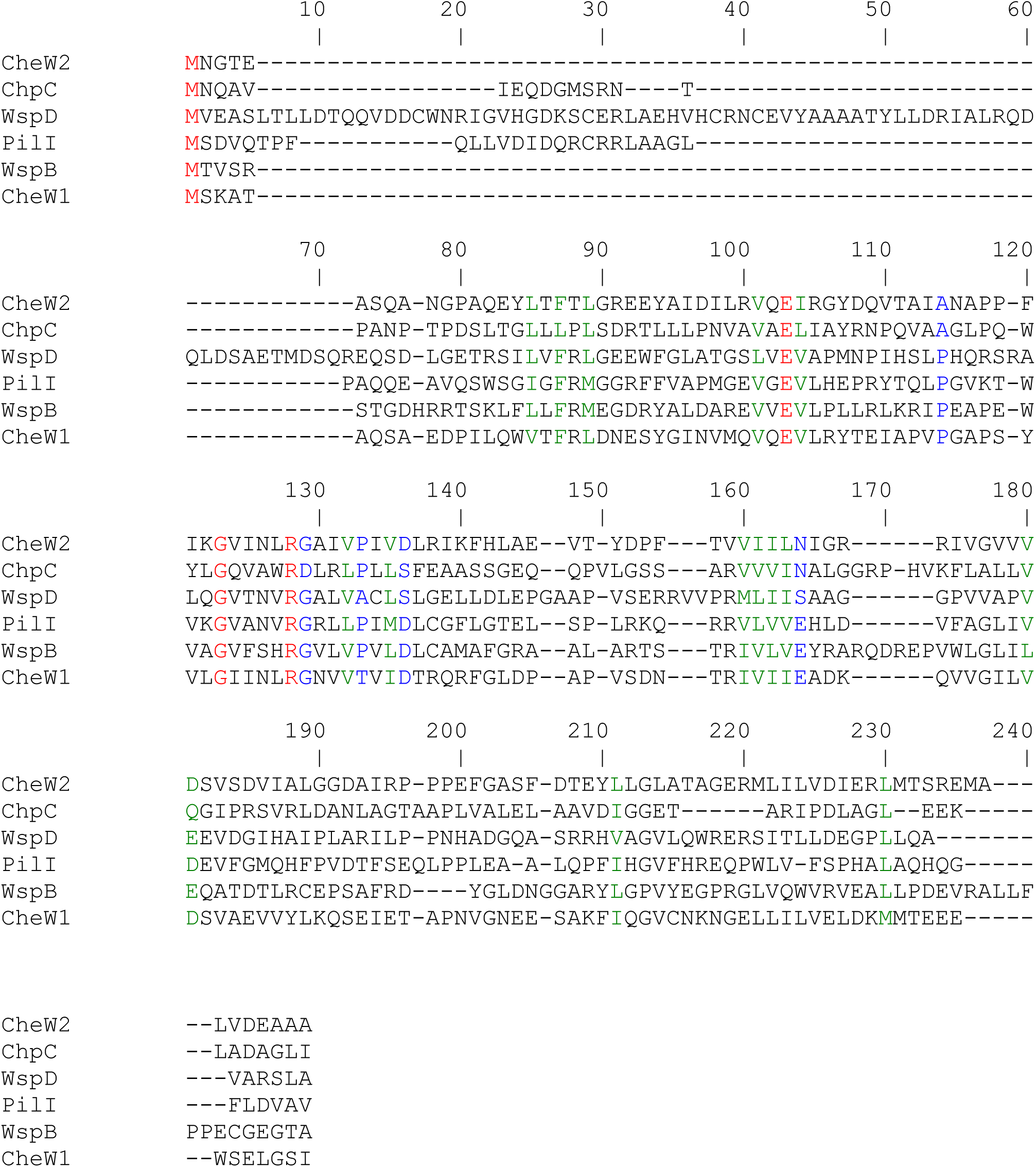
Sequence alignment of the CheW homologs of *Pseudomonas aeruginosa* PAO1. Sequences were aligned with the CLUSTALW algorithm of the npsa suite (*1*) using the GONNET weight matrix, a gap opening penalty of 10 and a gap extension penalty of 0.2. Red: identical; green: highly similar; blue: weakly similar.

**Fig. S2.**
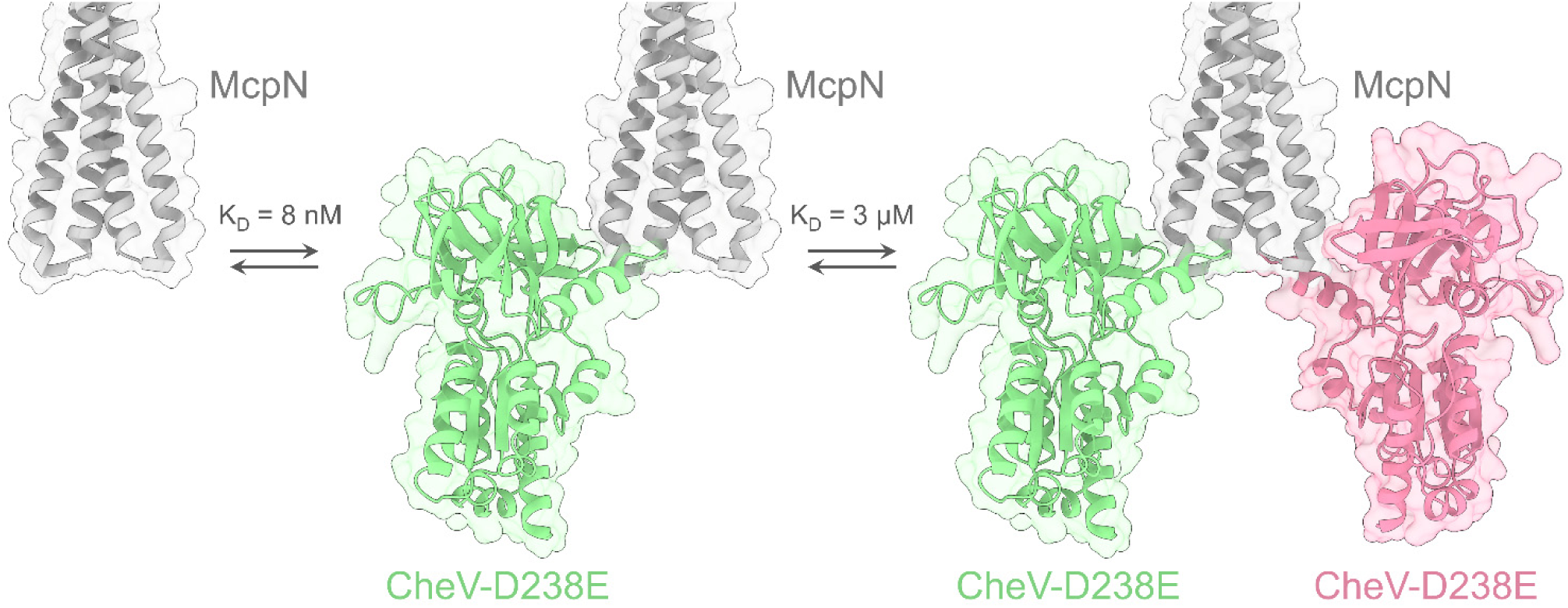
Hypothetical model of interaction of CheV D238E with the McpN signaling domain. This model is based on the isothermal titration calorimetry data shown in Fig. 5. Alphafold2 models of the dimeric McpN signaling domain and CheV D238E were generated and used for a structural alignment with the ternary complex chemoreceptor/CheW/CheA (domains P4 and P5) from *Thermotoga maritima* (PDB ID: 3UR1)(*2*). ITC data showed two binding events which are hypothesized to correspond to the sequential binding of two CheV to either side of the dimeric McpN signaling domain. The *K*_D_ values obtained by ITC are annotated.

**Fig. S3.**
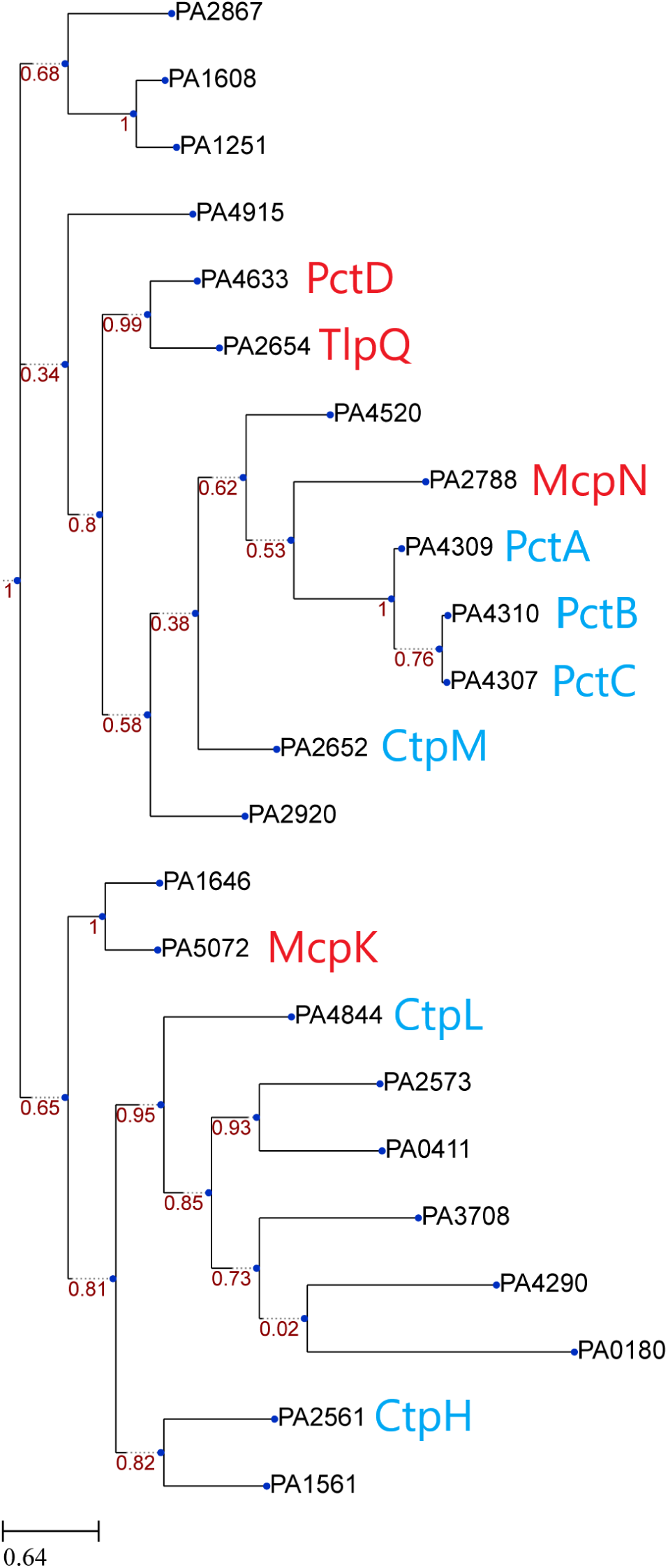
Sequence clustering of the cytosolic fragments of *P. aeruginosa* PAO1 transmembrane chemoreceptors. Chemoreceptors that were sensitive to the action of CheV are shown in red, and those that were insensitive in cyan (Fig. 3A). The cytosolic chemoreceptors PA1423 (BdlA), PA1930 (McpS) and PA0176 (McpB/Aer2) have not been included in this analysis. Analysis carried out using TREND and default settings (*3*).

**Fig. S4.**
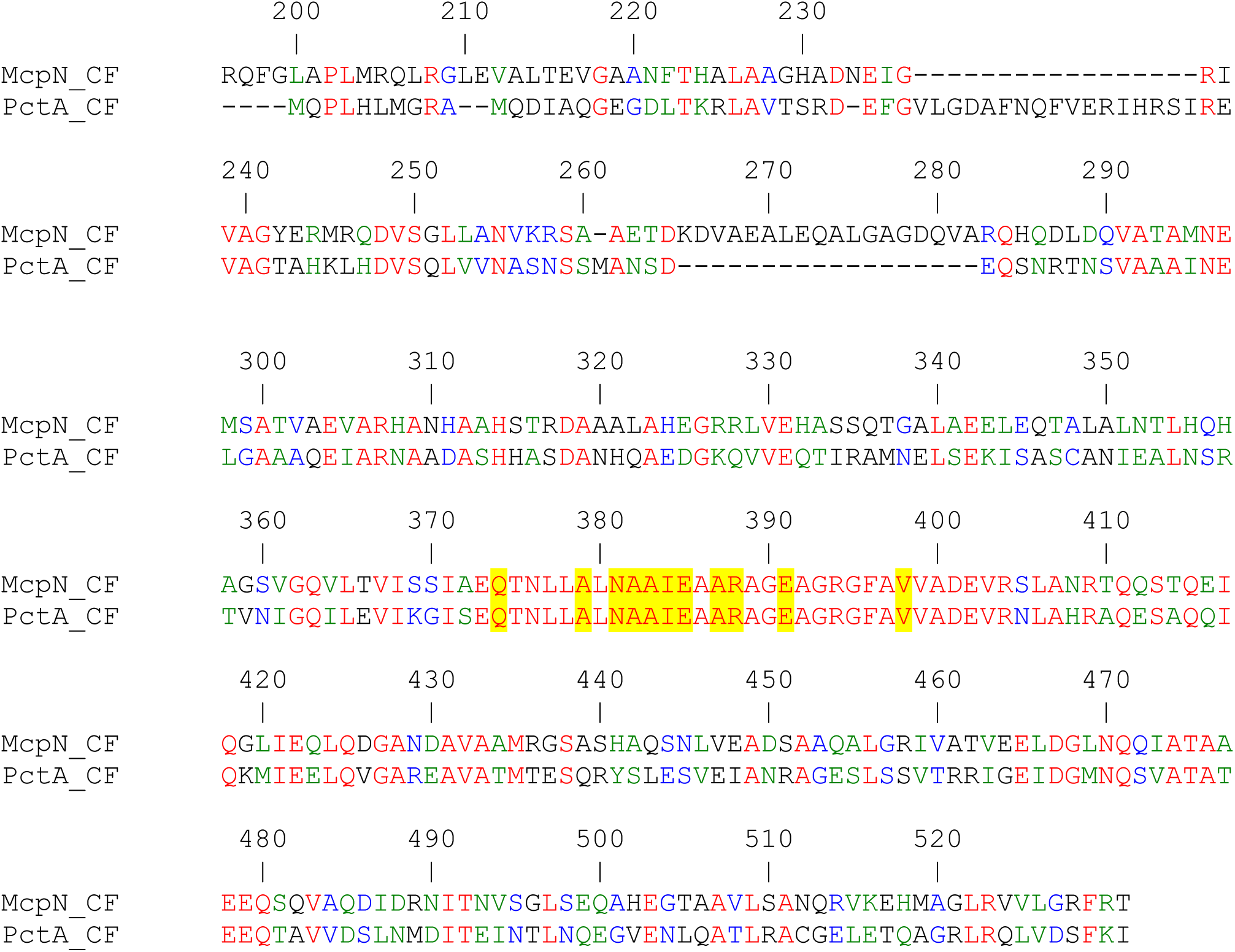
Sequence alignment of the cytosolic fragments of the McpN and PctA chemoreceptors of *P. aeruginosa* PAO1. Sequences were aligned with the CLUSTALW algorithm of the npsa suite (*1*) using the GONNET weight matrix, a gap opening penalty of 10 and a gap extension penalty of 0.2. Red: identical; green: highly similar; blue: weakly similar. The overall sequence identity is of 36 %. The positions of amino acids that were previously shown to interact with CheW (*4–6*) are shaded in yellow. McpN numbering is shown.

### Supplementary Tables

**Table S1.** Percentage of amino acids sequence identity derived from pairwise sequence alignments of the CheW proteins of *P. aeruginosa* PAO1. A multiple sequence alignment of these five proteins is shown in Fig. S1. Sequences were aligned with the CLUSTALW algorithm of the npsa suite (*1*) using the GONNET weight matrix, a gap opening penalty of 10 and a gap extension penalty of 0.2.

|  | **CheW<sub>1</sub>**<br>**(PA1464)** | **CheW<sub>2</sub>**<br>**(PA0177)** | **WspB**<br>**(PA3707)** | **WspD**<br>**(PA3705)** | **PilI**<br>**(PA0410)** | **ChpC**<br>**(PA0415)** |
| --- | --- | --- | --- | --- | --- | --- |
| **CheW<sub>1</sub>**<br>**(PA1464)** | X | 30 | 22 | 17 | 16 | 20 |
| **CheW<sub>2</sub>**<br>**(PA0177)** | 30 | X | 23 | 17 | 15 | 15 |
| **WspB**<br>**(PA3707)** | 22 | 23 | X | 14 | 19 | 19 |
| **WspD**<br>**(PA3705)** | 17 | 17 | 14 | X | 19 | 16 |
| **PilI**<br>**(PA0410)** | 16 | 15 | 19 | 19 | X | 18 |
| **ChpC**<br>**(PA0415)** | 20 | 15 | 19 | 16 | 18 | X |

**Table S2.** Relative protein quantification of *P. aeruginosa* PAO1 grown in different growth media. Shown are intensity-based absolute quantification (iBAQ) (x 10^6^) values of coupling proteins of PAO1 grown in minimal medium (MM) supplemented with 0.2 mM inorganic phosphate (Pi), MM supplemented with 1 mM Pi and LB medium. IBAQ values permit a comparison of protein levels. Data have been extracted from (*7*).

|  | **MM + 0.2 mM Pi**<br>**(ratio to CheV)** | **MM + 1 mM Pi**<br>**(ratio to CheV)** | **LB**<br>**(ratio to CheV)** |
| --- | --- | --- | --- |
| **CheV (PA3349)** | 34.25 (1) | 24.22 (1) | 17.01 (1) |
| **CheW<sub>1</sub> (PA1464)** | 61.32 (1.79) | 51.56 (2.13) | 70.93 (4.17) |
| **CheW<sub>2</sub> (PA0177)** | 1.26 (0.37) | 0.11 (0.005) | Not detected |
| **WspB (PA3707)** | 0.19 (0.005) | 0.31 (0.01) | 0.30 (0.02) |
| **WspD (PA3705)** | 0.97 (0.03) | 0.29 (0.01) | 0.27 (0.02) |
| **PilI (PA0410)** | 2.49 (0.07) | 2.82 (0.11) | 6.67 (0.39) |
| **ChpC (PA0415)** | 3.96 (0.11) | 3.43 (0.14) | 6.86 (0.40) |

**Table S3.**
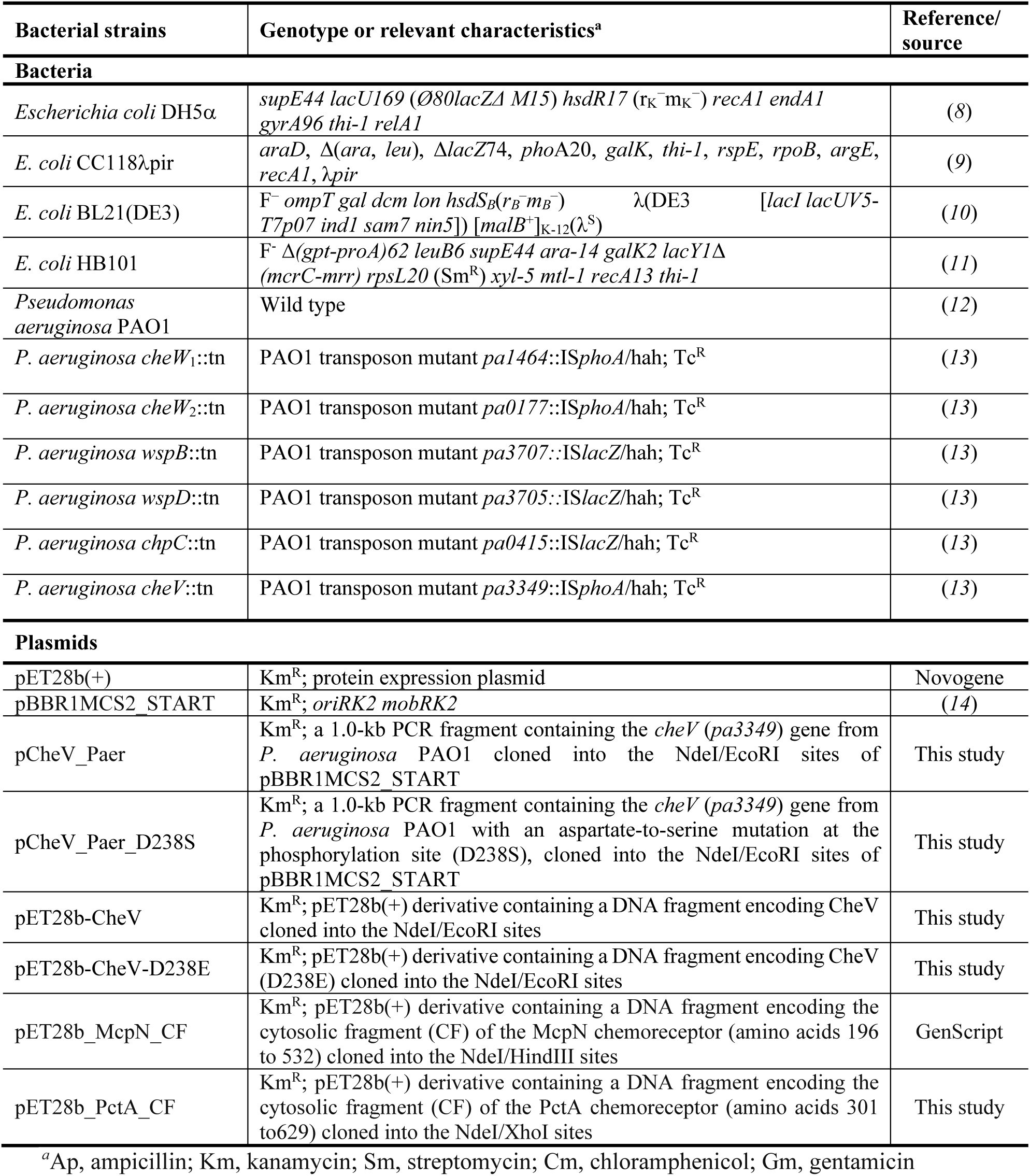
Bacteria and plasmids used in this study.

**Table S4.**
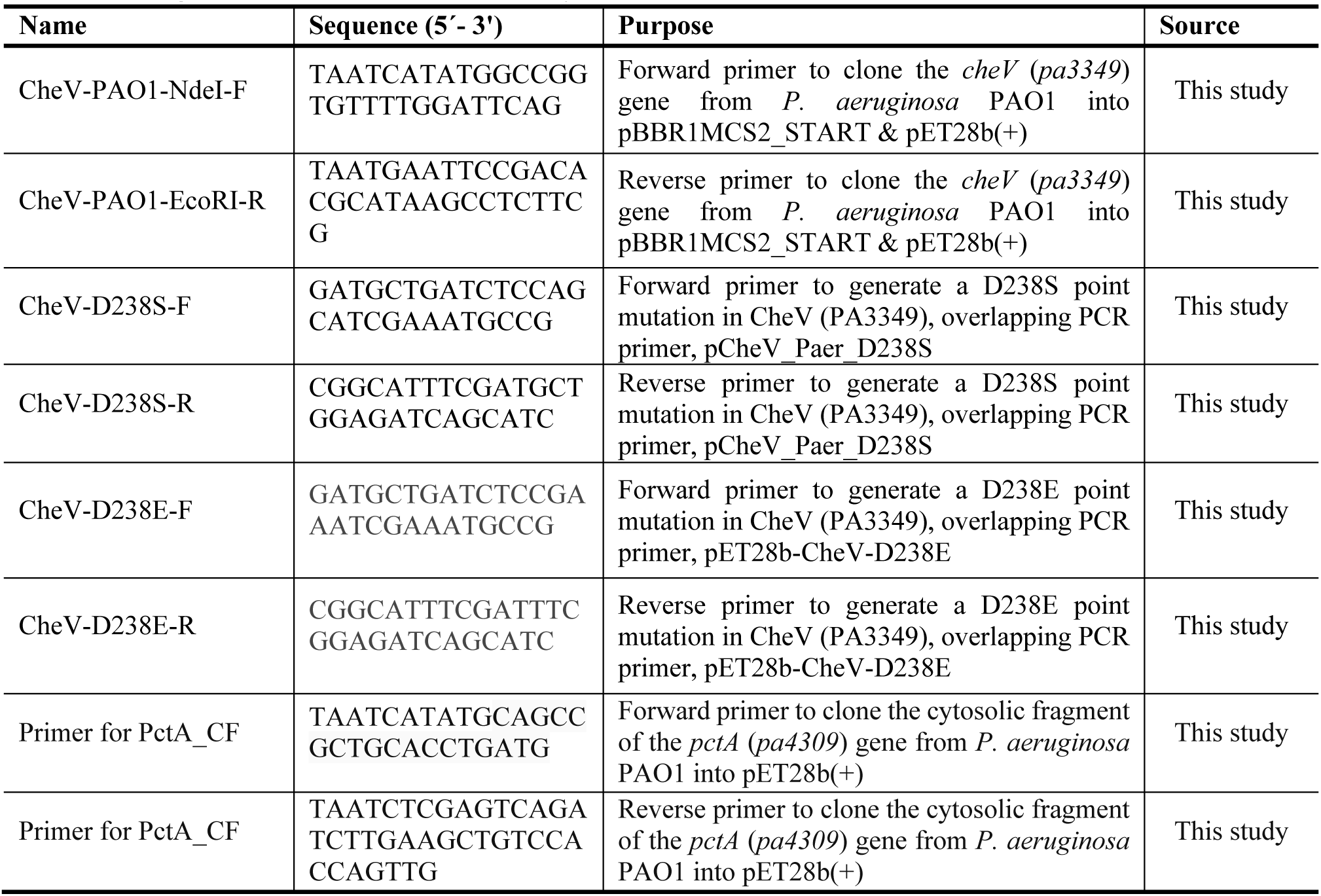
Oligonucleotides used in this study.

